# Cardiac *Nmnat*/NAD^+^/SIR2 pathways activation mediates endurance exercise resistance to heart defects induced by a high-fat diet in aging *Drosophila*

**DOI:** 10.1101/2021.02.10.430616

**Authors:** Deng-tai Wen, Lan Zheng, Kai Lu, Wen-qi Hou

## Abstract

Endurance exercise is an important way to resist and treat a high-fat-diet(HFD)-induced heart defects, but the underlying molecular mechanisms are poorly understood. Here, we used *Drosophila* to identify whether cardiac Nmnat/NAD^+^/SIR2 pathways activation could mediate endurance exercise resistance to heart defects. The results showed that endurance exercise activated the cardiac Nmnat/NAD^+^/SIR2/*FOXO* pathway and Nmnat/NAD^+^/SIR2/*PGC-1α* pathway, including up-regulating cardiac *Nmnat, SIR2, FOXO, PGC-1α* expression, SOD activity, and NAD^+^ level, and it prevented HFD-induced or cardiac *Nmnat* knock-down-induced cardiac lipid accumulation, MDA content and fibrillation increase, and fractional shortening decrease. Cardiac *Nmnat* overexpression activated heart Nmnat/NAD^+^/SIR2 pathways and resisted HFD-induced cardiac malfunction, but it could not protect against HFD-induced lifespan reduction and locomotor impairment. Exercise improved lifespan and mobility in cardiac *Nmnat* knockdown flies. Therefore, current results confirmed that cardiac Nmnat/NAD^+^/SIR2 pathways were important antagonists of HFD-induced heart defects. The cardiac *Nmnat/NAD^+^/SIR2* pathways activation was the important underlying molecular mechanism of endurance exercise and cardiac Nmnat overexpression against heart defects in *Drosophila.*

## Introduction

Heart disease is the number one cause of human death in contemporary society. Many reports have shown the strong connections between obesity and cardiac dysfunction in both animals and humans^1,2^. The epidemic of obesity and its associated cardiac dysfunctions are at least partly caused by consumption of high caloric fat- and sugar-enriched foods^3^. Because of genetically and metabolically complex, the biology and physiology of cardiac dysfunctions induced by a High-fat-diet (HFD) remain poor understanding in humans. Among the invertebrate models, *Drosophila* is the only one with a heart, and it has been a good genetic tool for analyzing heart function. Importantly, since most gene families and signaling pathways are highly conserved between flies and human, flies can be used to perform heart function-related studies well^1^. For example, recent studies show that consumption of high-fat diets in *Drosophila* leads to excessive fat accumulation accompanied by severe heart defects, including increased frequency of arrhythmias, reduced cardiac output, increased non-contractile myocardial cells, and altered myofibrillar structure and collagen content^4–7^. Moreover, it has been identified that cardiac *PGC-1α*, *FOXO,* and *bmm* are key antagonists of HFD-induced heart defects in flies ^5,8^. Therefore, compared with mammals, the regulatory network of heart defects is easier to study in *Drosophila.*

In both mammals and *Drosophila,* physical exercise is regard as a cost-effective approach to prevent or improve some heart diseases. For example, exercise can mitigate HFD-induced cardiac fibrosis, fractional shortening reduction, triacylglycerol(TAG) accumulation, and arrhythmias in both rats and flies^4,9,10^. However, since both heart defects and cardiac exercise adaptation are associated with complex molecular mechanisms, the regulatory network of exercise resistance heart defects still remains poor understanding. Increasing evidence suggests that the moderate exercise may be an upstream regulator of PGC-1α, FOXO, SIRT1, and NAD^+11^. For instance, endurance exercise can protect myocardium by reducing the myocardial oxidative stress injury and apoptosis via activating SIRT1 signaling pathway, up-regulating the myocardial expression of SIRT1 and regulating the deacetylation of FOXO in rats^12^. Besides, the reduction of PGC-1α in the heart is insufficient to cause an aging phenotype, and moderate overexpression of *PGC-1α* reduces pathological remodeling of older hearts and contributes to the beneficial effects of exercise on cardiac function in aging ^13^. Next, endurance exercise results in a sustained increase in NAD^+^ levels in gastrocnemius muscle of rats ^14^. Finally, our previous research suggests that endurance exercise can improve heart dysfunction induced by CG9940 knock-down ^15^, which may be related to the up-regulation of NAD^+^ concentration in heart by exercise. Therefore, the cardiac NAD^+^/SIR2/PGC-1α and NAD^+^/SIR2/FOXO may be two key pathways for exercise to combat lipotoxic cardiomyopath induced by a HFD. However, no systematic studies have confirmed this speculation.

Nicotinamide mononucleotide adenylyltransferase (Nmnat) is initially identified as an NAD^+^ synthase. It catalyzes the reversible conversion of NMN (nicotinamide mononucleotide) to NAD^+^ in the final step of both the de novo biosynthesis and salvage pathways in most organisms across all three kingdoms of life. Nicotinamide adenine dinucleotide (NAD) is an essential co-cofactorthat servesto mediate various biological processes, including metabolism, DNA repair, and gene expression^16,17^. Increasing evidence shows that Nmnat is indispensable in maintaining neuronal homeostasis, for example, *Nmnat* is closely related to AD and other tauopathies^18^. The NMNAT is a rate-limiting enzyme present in all organisms, and it reversibly catalyzes the important step in the biosynthesis of NAD from ATP and NMN ^19^. Overexpression of *Nmnat* has been reported can increase NAD^+^ levels in cells^20,21^. However, the function of *Nmnat* gene in the heart is still unknown.

To explore whether endurance exercise or cardiac *Nmnat* overexpression can resist HFD-induced heart defects via activating NAD^+^/SIR2 pathways, fruit flies will be applied in this study. Firstly, flies were taken endurance exercise or fed a HFD to explore whether these two interventions can change the activity of cardiac Nmnat/NAD^+^/SIR2 pathways. Next, cardiac *Nmnat* overexpression line was built by UAS/hand-Gal4 system, and these flies were fed a HFD to explore whether up-regulation of cardiac NAD^+^/SIR2 pathways can protect heart from lipotoxic cardiomyopath induced by a HFD. Finally, the cardiac *Nmnat* knock-down line was constructed by RNAi to explore whether down-regulation of cardiac NAD^+^/SIR2 pathways can induce lipotoxic cardiomyopath. The cardiac *Nmnat* knock-down flies were taken endurance exercise to explore whether endurance exercise can improve lipotoxic cardiomyopath induced by cardiac *Nmnat* knock-down. The climbing index and lifespan were also measured in cardiac *Nmnat* overexpression and RNAi flies to know the relationship between exercise, a HFD, lipotoxic cardiomyopath, and health-span.

## Results

### Exercise prevents HFD-induced cardiac dysfunction and Nmnat/NAD^+^/SIR2 pathways suppression

Increasing study reports that a HFD causes cardiac dysfunction in both animals and flies, such as increased frequency of arrhythmias, reduced cardiac output, increased fibrillations and TAG levels. Endurance exercise has been reports can protect heart from HFD-induced cardiac malfunction, but its molecular mechanisms of regulation are still poorly understood. Accumulating evidence suggests that endurance exercise, HFD-induced cardiac malfunction, and Nmnat/NAD^+^/SIR2 pathways are closely related. So, to explore whether endurance exercise resistance HFD-induced lipotoxic cardiomyopath was accompanied by activating cardiac NAD^+^/SIR2 pathways, flies were taken endurance exercise or fed a HFD in this study. The cardiac TAG levels and *bmm* expression were measured to show the status of lipid accumulation of heart. The cardiac *Nmnat* expression, NAD^+^ levels, *SIR2* expression, *FOXO* expression, *PGC-1α* expression, SOD activity levels, and MDA levels were measured to reflect the activity of cardiac Nmnat/NAD^+^/SIR2 pathways. The heart rate, fractional shortening, diastolic diameter, systolic diameter, and fibrillation were tested by M-mode trace to represent the status of heart contractility and dysfunction.

We found that exercise significantly decreased cardiac TAG content (2-factor ANOVA, P < 0.01), and HFD significantly increased cardiac TAG content (2-factor ANOVA, P < 0.01). Exercise and HFD had no interaction influence on TAG content (2-factor ANOVA, P > 0.05). Besides, we also found that exercise significantly increased cardiac *bmm* expression (2-factor ANOVA, P < 0.01), and HFD significantly reduced cardiac *bmm* expression (2-factor ANOVA, P < 0.01). Exercise and HFD had no interaction influence on cardiac *bmm* expression (2-factor ANOVA, P > 0.05). Further analysis and comparison of results showed that the relative heart TAG content in Nmnat-C+E group flies was lower than Nmnat-C group flies (LSD test, P < 0.05); the relative heart TAG content in Nmnat-C+HFD group flies was higher than Nmnat-C group flies (LSD test, P < 0.01); the relative heart TAG content in Nmnat-C+HFD+E group flies was lower than Nmnat-C+HFD group flies (LSD test, P < 0.01), and there was no significant difference between Nmnat-C group flies and Nmnat-C+HFD+E group flies in the relative TAG content(LSD test, P > 0.05) (**Fig.1-a**). Moreover, the *Brummer (bmm)* encodes the *Drosophila* homologue of mammalian ATGL lipase, and it also mediates fat hydrolysis in this invertebrate model system^22^. A HFD can reduce *bmm* expression, and *bmm* overexpression efficiently prevents HFD-induced fat accumulation and heart dysfunction^5^. The results showed that the relative heart *bmm* expression in Nmnat-C+E group flies was higher than Nmnat-C group flies (LSD test, P < 0.01); the relative heart *bmm* expression in Nmnat-C+HFD group flies was lower than Nmnat-C group flies (LSD test, P < 0.01); the relative heart *bmm* expression in Nmnat-C+HFD+E group flies was higher than Nmnat-C+HFD group flies (LSD test, P < 0.01), and the relative heart *bmm* expression in Nmnat-C+HFD+E group flies was higher than Nmnat-C group flies (LSD test, P < 0.05) (**Fig.1-b**). These results suggested that endurance exercise resisted HFD-induced heart lipid accumulation by promoting the breakdown of TAG.

**Fig.1.**
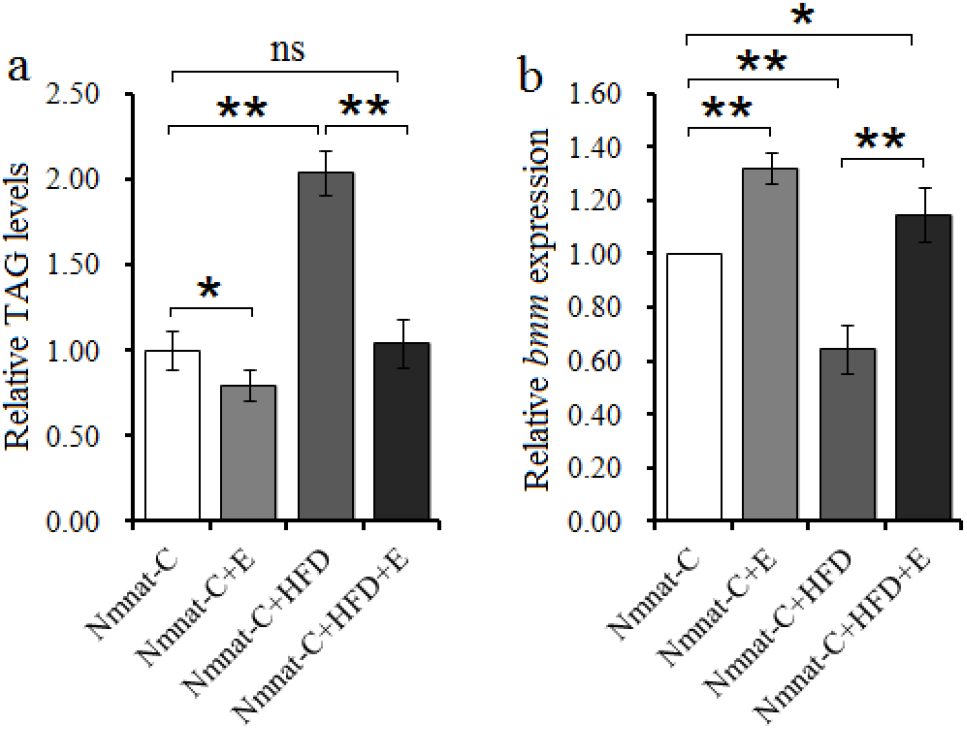
The effect of a HFD and exercise on heart lipid accumulation. **(a)** Heart relative TAG level. Results are expressed as the fold difference compared with Nmnat-C flies. The sample size was 80 hearts with 3 biological replicates. **(b)** Relative heart *bmm* gene expression level. The sample size was 80 hearts with 3 biological replicates. A 2-way ANOVA was used to identify differences among the Nmnat-C, Nmnat-C+E, Nmnat-C+HFD, and Nmnat-C+HFD+E groups flies. Data are represented as means ± SEM. \**P* <0.05; \*\**P* <0.01

For cardiac Nmnat/NAD^+^/SIR2 pathways, the results showed that exercise significantly increased cardiac Nmnat expression, NAD^+^ level, *SIR2* expression, *FOXO* expression, *PGC-1α* expression, and SOD activity level, and it notably reduced cardiac MDA level (2-factor ANOVA, P < 0.01). On the contrary, a HFD significantly increased cardiac MDA levels, and it notably decreased cardiac Nmnat expression, NAD^+^ levels, *SIR2* expression, *FOXO* expression, *PGC-1α* expression, and SOD activity level (2-factor ANOVA, P < 0.01). Exercise and a HFD had no interaction influence on cardiac *Nmnat* expression, NAD^+^ level, *SIR2* expression, *FOXO* expression, *PGC-1α* expression, SOD activity level, and MDA level (2-factor ANOVA, P > 0.05). Further analysis and comparison of results showed that the relative cardiac *Nmnat* expression, NAD^+^ level, *SIR2* expression, *FOXO* expression, *PGC-1α* expression, and SOD activity level in Nmnat-C+E group flies were higher than Nmnat-C group flies (LSD test, P < 0.01 or P < 0.05), but the cardiac MDA level in Nmnat-C+E group flies was lower than Nmnat-C group flies (LSD test, P < 0.05); besides, the cardiac MDA level in Nmnat-C+HFD group flies was higher than Nmnat-C group flies (LSD test, P < 0.01), but the relative cardiac *Nmnat* expression, NAD^+^ level, *SIR2* expression, *FOXO* expression, *PGC-1α* expression, and SOD activity level in Nmnat-C+HFD group flies were lower than Nmnat-C group flies (LSD test, P < 0.01 or P < 0.05); Moreover, the relative cardiac MDA level in Nmnat-C+HFD+E group flies was lower than Nmnat-C+HFD group flies, and the relative cardiac *Nmnat* expression, NAD^+^ level, *SIR2* expression, *FOXO* expression, *PGC-1α* expression, and SOD activity level in Nmnat-C+HFD+E group flies was higher than Nmnat-C+HFD group flies (LSD test, P < 0.01); finally, there was no significant difference between Nmnat-C group flies and Nmnat-C+HFD+E group flies in the cardiac *Nmnat* expression, NAD^+^ level, *SIR2* expression, *FOXO* expression, MDA level, and SOD activity level, but the relative heart *PGC-1α* expression in Nmnat-C+HFD+E group flies was higher than Nmnat-C group flies (LSD test, P < 0.05) (**Fig.2-a, b, c, d, e, f, and g**). These results indicated that endurance exercise relieved HFD-induced lipid toxicity injury in myocardial cells by up regulating Nmnat/NAD^+^/SIR2/FOXO pathway and Nmnat/NAD^+^/SIR2/*PGC-l*« pathway.

**Fig.2.**
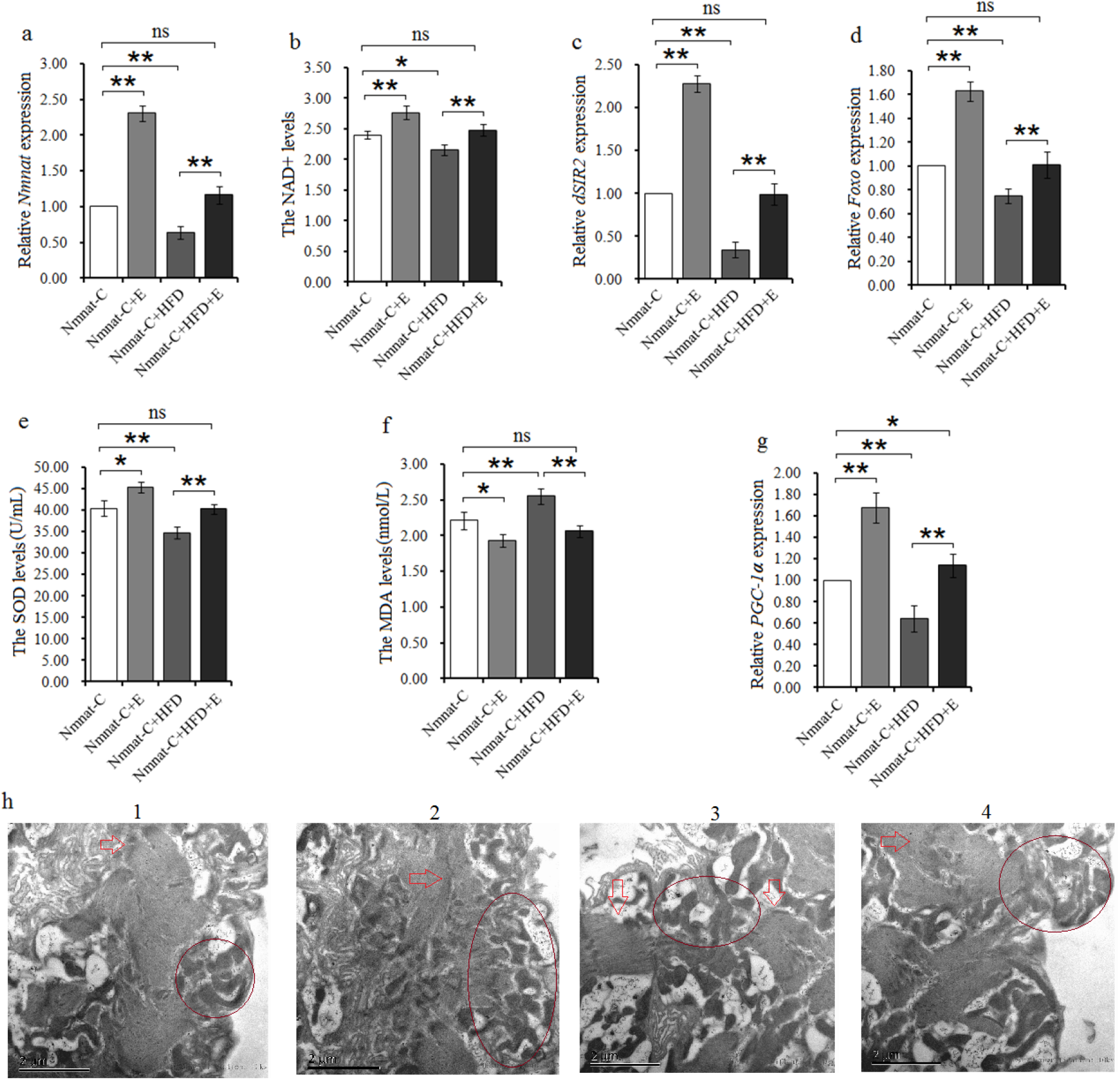
The effect of a HFD and exercise on Nmnat/NAD^+^/SIR2 pathways. **(a)** Heart relative *Nmnat* expression. **(b)** Heart NAD^+^ levels. **(c)** Heart relative *SIR2* expression. **(d)** Heart relative *FOXO* expression. **(e)** Heart relative SOD activity levels. **(f)** Heart relative MDA levels. **(g)** Heart relative *PGC-1α* expression. **(h)**The images of transmission electron microscopy; 1: *Nmnat-C;* 2: *Nmnat-C+E;* 3: *Nmnat*-C+HFD; 4: *Nmnat*-C+HFD+E. The pictures showed endurance exercise increased the mitochondrial numbers and improved myofibril arrangement regularity in myocardial cells. The circle pointed the position of the mitochondria in myocardial cells. The rectangle pointed the position of the Z line in myocardial cells. A 2-way ANOVA was used to identify differences among the Nmnat-C, Nmnat-C+E, Nmnat-C+HFD, and Nmnat-C+HFD+E groups flies. Data are represented as means ± SEM. \**P <0.05; **P <0.01.* The sample size of each indicator was 80 hearts, with 3 biological replicates.

For cardiac function, the results showed that exercise significantly increased cardiac fractional shortening, and it notably reduced heart rate and fibrillation (2-factor ANOVA, P < 0.05 or P < 0.01). On the contrary, a HFD significantly increased heart rate and fibrillation (2-factor ANOVA, P < 0.05 or P < 0.01), but it notably reduced fractional shortening (2-factor ANOVA, P < 0.05). Exercise and a HFD had no interaction influence on cardiac fractional shortening, heart rate, and fibrillation (2-factor ANOVA, P > 0.05). Both exercise and a HFD also had no significant influence on diastolic diameter and systolic diameter, and they had no interaction influence on diastolic diameter and systolic diameter (2-factor ANOVA, P > 0.05). Further analysis and comparison of results showed that the fractional shortening in Nmnat-C+E group flies was higher than Nmnat-C group flies (LSD test, P < 0.05), but the heart rate and fibrillation in Nmnat-C+E group flies were lower than Nmnat-C group flies (LSD test, P < 0.05 or P < 0.01); in addition, the heart rate and fibrillation in Nmnat-C+HFD group flies were higher than Nmnat-C group flies (LSD test, P < 0.05 or P < 0.01), but the fractional shortening in Nmnat-C+HFD group flies was lower than Nmnat-C group flies (LSD test, P < 0.05); next, the heart rate and fibrillation in Nmnat-C+HFD+E group flies were lower than Nmnat-C+HFD group flies, and the fractional shortening in Nmnat-C+HFD+E group flies was higher than Nmnat-C+HFD group flies (LSD test, P < 0.01); finally, there was no significant difference between Nmnat-C group flies and Nmnat-C+HFD+E group flies in the heart rate, fractional shortening, and fibrillation(LSD test, P > 0.05)(**Fig.3-a, b, c, d, e, f, and g**). These results indicated that endurance exercise improved HFD-induced heart weak contractility and severe arrhythmia.

**Fig.3.**
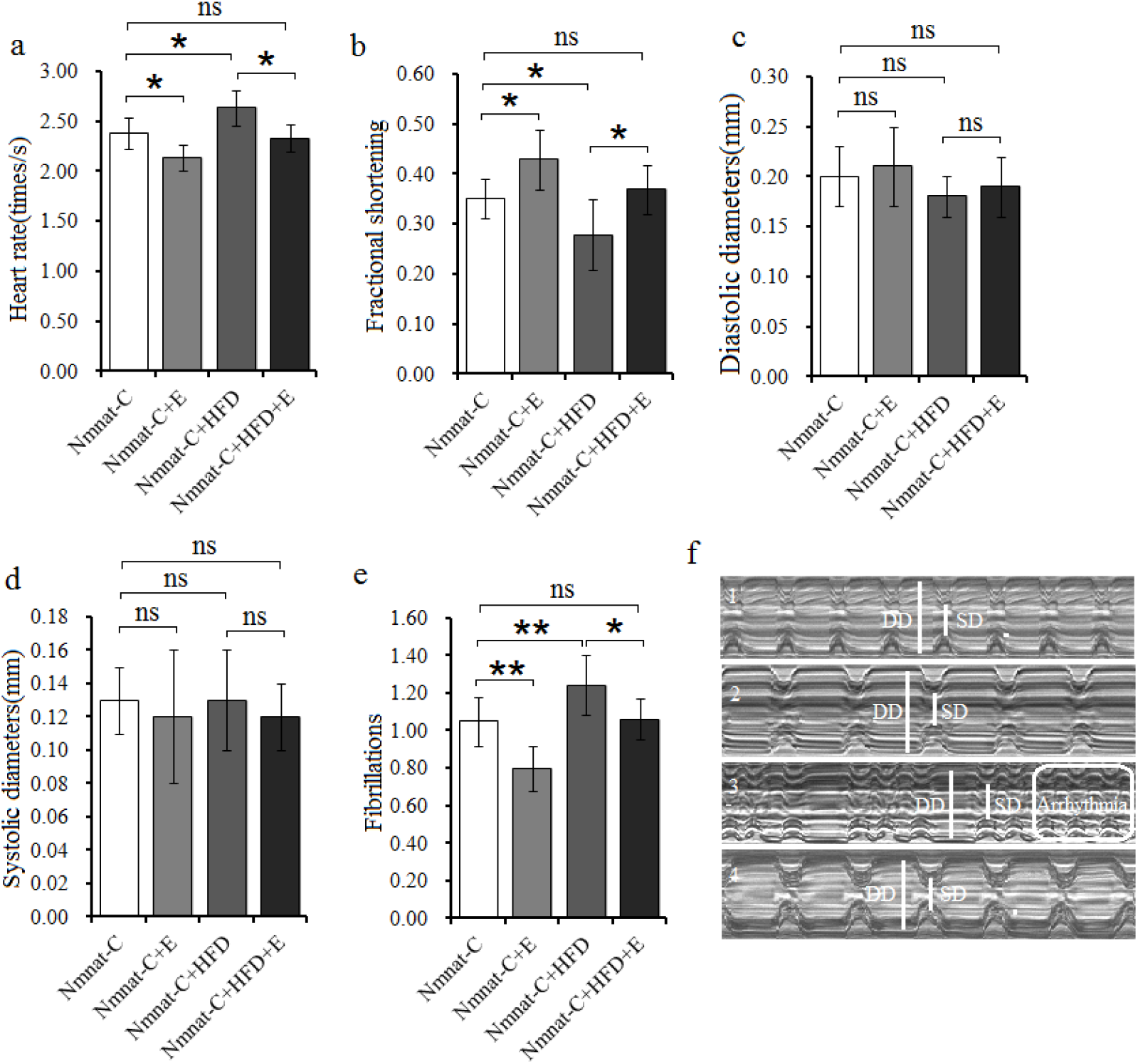
The effect of a HFD and exercise on heart functions. **(a)** Heart rate. **(b)** Fractional shortening. **(c)** Cardiac diastolic diameter. **(d)** Cardiac systolic diameter. **(e)** Fibrillations. **(f)** Illustrating qualitative differences in heart function parameters (3s): fractional shortening and arrhythmia index; 1: *Nmnat-C;* 2: *Nmnat-C+E;* 3: *Nmnat*-C+HFD; 4: *Nmnat*-C+HFD+E. A 2-way ANOVA was used to identify differences among the Nmnat-C, Nmnat-C+E, Nmnat-C+HFD, and Nmnat-C+HFD+E groups flies. Data are represented as means ± SEM. *P <0.05; **P <0.01. The sample size was 30 hearts.

### Exercise improves *Nmnat* RNAi-induced heart dysfunction and lipid accumulation

Functional studies in *Drosophila* and mammals have shown that loss of *Nmnat* causes neurodegeneration and a decreased in NAD^+^ levels of cells ^20,21^. Although previous results indicated that exercise prevention HFD-induced cardiac dysfunction was accompanied by up-regulating cardiac Nmnat/NAD^+^/SIR2 pathways activity, it still remained unclear whether cardiac Nmnat/NAD^+^/SIR2 pathways could mediate endurance exercise resist HFD-induced cardiac malfunction since there was no direct evidence that cardiac Nmnat/NAD^+^/SIR2 pathways were the key pathways for HFD-induced lipid toxicity in cardiomyopathy. Therefore, to explore whether cardiac Nmnat/NAD^+^/SIR2 pathways could modulate the formation of HFD-induced lipotoxic cardiomyopath, the cardiac *Nmnat* gene was knocked down by RNAi.

The results showed that cardiac relative *Nmnat* expression in Nmnat-KD group flies was lower than Nmnat-C group flies (LSD test, P < 0.01), which suggested that the cardiac *Nmnat* was successfully knocked down by RNAi (**Fig.5-a**). Besides, for cardiac Nmnat/NAD^+^/SIR2 pathways, the results showed that the cardiac NAD^+^ level, *SIR2* expression, FOXO expression, SOD activity level, and *PGC-1α* expression in Nmnat-KD group flies were lower than Nmnat-C group flies (LSD test, P < 0.01) (**Fig.5-b, c, d, e, and g**), but the MDA level in Nmnat-KD group flies was higher than Nmnat-C group flies (LSD test, P < 0.01) (**Fig.5-f**). These results suggested that down-regulation of cardiac *Nmnat* expression reduced the activity of cardiac Nmnat/NAD^+^/SIR2 pathways and increased the risk of oxidative damage to myocardial cells. Moreover, the cardiac TAG level in Nmnat-KD group flies was higher than Nmnat-C group flies (LSD test, P < 0.01), and the cardiac *bmm* expression in Nmnat-KD group flies was lower than Nmnat-C group flies (**Fig.5-h and i**). These results suggested that down-regulation of cardiac *Nmnat* expression increased heart lipid accumulation. Finally, the heart rate, diastolic diameter, systolic diameter, and fibrillation in Nmnat-KD group flies were higher than Nmnat-C group flies (LSD test, P < 0.05 or P < 0.01), and the fractional shortening in Nmnat-KD group flies were lower than Nmnat-C group flies (LSD test, P < 0.05) (**Fig.5-j, k, l, m, n, and o**). These results indicated that down-regulation of cardiac *Nmnat* expression weakened the heart contractility and exacerbated cardiac arrhythmia. Therefore, we claimed that cardiac *Nmnat* knock-down could induce lipotoxic cardiomyopath by reducing cardiac Nmnat/NAD^+^/SIR2 pathways.

**Fig.5.**
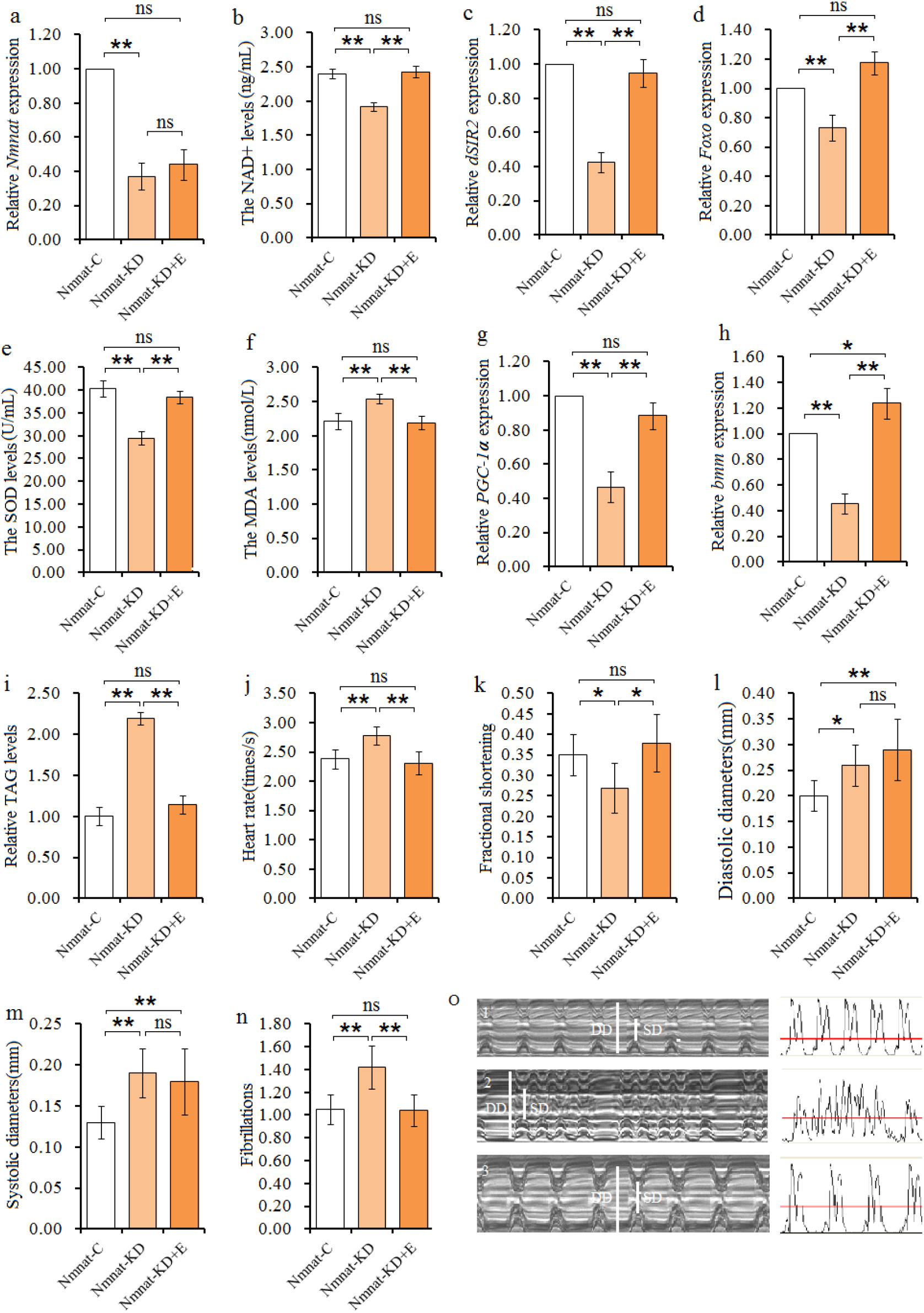
The effect of cardiac *Nmnat* RNAi and endurance exercise on heart. **(a)** Heart relative *Nmnat* expression. **(b)** Heart NAD^+^ level. **(c)** Heart relative *SIR2* expression. **(d)** Heart relative *FOXO* expression. **(e)** Heart relative SOD activity levels. **(f)** Heart relative MDA levels. **(g)** Heart relative *PGC-1α* expression. **(h)** Heart relative *bmm* gene expression level. **(i)** Heart relative TAG level. **(j)** Heart rate. **(k)** Fractional shortening. **(l)** Cardiac diastolic diameter. **(m)** Cardiac systolic diameter. **(n)** Fibrillations. **(o)** Illustrating qualitative differences in heart function parameters (3s): fractional shortening and arrhythmia index; 1: *Nmnat-C;* 2: *Nmnat-KD;* 3: *Nmnat-KD+E;* Data are represented as means ± SEM. *P <0.05; **P <0.01.

Next, to further explore the relationship between endurance exercise and cardiac Nmnat/NAD^+^/SIR2 pathways, the cardiac *Nmnat* knock-down flies were taken endurance exercise. The results showed that the cardiac NAD^+^ level, *SIR2* expression, FOXO expression, SOD activity level, and *PGC-1α* expression in Nmnat-KD group flies were higher than Nmnat-KD+E group flies (LSD test, P < 0.01) (**Fig.5-a, b, c, d, e, and g**), but the MDA level in Nmnat-KD+E group flies was lower than Nmnat-KD group flies (LSD test, P < 0.01) (**Fig.6-f**). However, there was no significant difference in cardiac relative *Nmnat* expression between Nmnat-C flies and Nmnat-KD+E flies (LSD test, P > 0.05), which indicated exercise training did not change *Nmnat* expression in Nmnat-RNAi flies. These results indicated that endurance exercise improved cardiac NAD^+^/SIR2 pathways activity by other pathways of NAD synthesis other than the Nmnat pathway, and exercise decreased the risk of oxidative damage to myocardial cells. In addition, the cardiac TAG level in Nmnat-KD+E group flies was lower than Nmnat-KD group flies (LSD test, P < 0.01), and the cardiac *bmm* expression in Nmnat-KD+E group flies was higher than Nmnat-KD group flies (**Fig.5-h and i**). These results indicated that endurance exercise accelerated TAG catabolism and prevented heart lipid accumulation. Finally, the results displayed that the heart rate and fibrillation in Nmnat-KD+E group flies were lower than Nmnat-KD group flies (LSD test, P < 0.05), and the fractional shortening in Nmnat-KD+E group flies were higher than Nmnat-KD group flies (LSD test, P < 0.05) (**Fig.5-j, k, n, and o**). These results indicated that exercise improved heart contractility defect and cardiac arrhythmia induced by cardiac *Nmnat* knock-down. So, we claimed that endurance exercise could improve lipotoxic cardiomyopath induced by cardiac *Nmnat* knock-down by activating cardiac NAD^+^/SIR2 pathways.

**Fig.6.**
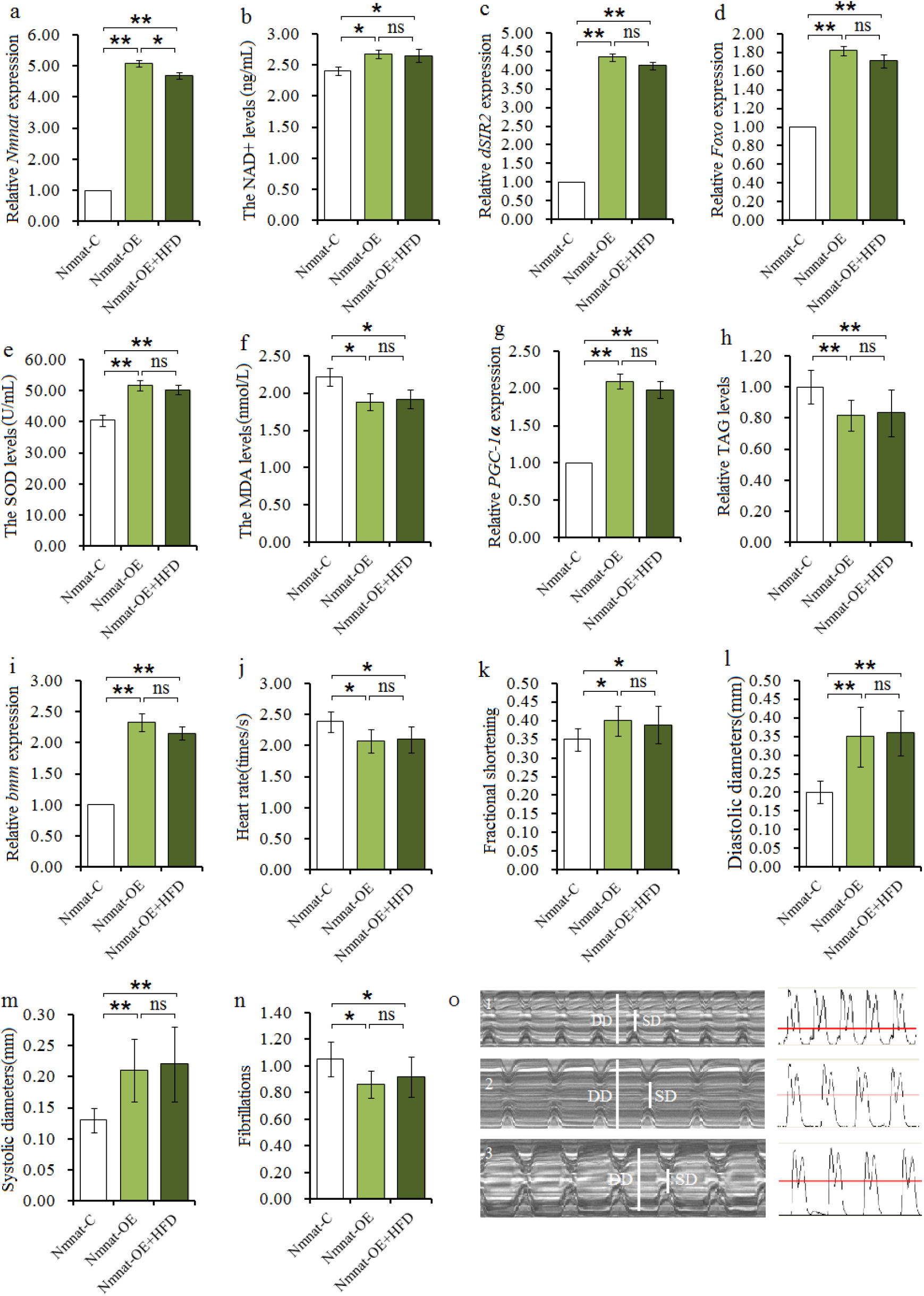
The effect of cardiac *Nmnat* overexpression and HFD on heart. **(a)** Heart relative *Nmnat* expression. **(b)** Heart NAD^+^ level. **(c)** Heart relative *SIR2* expression. **(d)** Heart relative *FOXO* expression. **(e)** Heart relative SOD activity levels. **(f)** Heart relative MDA levels. **(g)** Heart relative *PGC-1α* expression. **(h)** Heart relative *bmm* gene expression level. **(i)** Heart relative TAG level. **(j)** Heart rate. **(k)** Fractional shortening. **(l)** Cardiac diastolic diameter. **(m)** Cardiac systolic diameter. **(n)** Fibrillations. **(o)** Illustrating qualitative differences in heart function parameters (3s): fractional shortening and arrhythmia index; 1: *Nmnat-C;* 2: *Nmnat-OE;* 3: *Nmnat-* OE+HFD; Data are represented as means ± SEM. *P <0.05; **P <0.01.

To understand the improvement degree of endurance exercise on heart defects induced by cardiac *Nmnat* knockdown, we compared Nmnat-C group flies with Nmnat-KD+E group flies. The results showed that there was no significant difference in cardiac relative *Nmnat* expression, NAD^+^ level, *SIR2* expression, FOXO expression, SOD activity level, *PGC-1α* expression, TAG level, heart rate, fractional shortening, and fibrillation between Nmnat-C flies Nmnat-KD+E flies (LSD test, P > 0.05) (**Fig.5-a, b, c, d, e, g, i, j, k, and o**). These results indicated that endurance exercise could almost reverse lipotoxic cardiomyopath induced by cardiac *Nmnat* knock-down and return the heart to the statue of cardiac *Nmnat* normal expression.

### Cardiac *Nmnat* overexpression resists HFD-induced lipotoxic cardiomyopath

It has been reported that overexpression of *Nmnat* could delay age-related neurodegeneration, and it also suppresses dendrite maintenance defects associated with loss of the tumor suppressor kinase Warts^23^. Besides, *Nmnat* overexpression can improve health-span and locomotor activity in aging *Drosophila* ^24^. Although our previous results showed that loss of *Nmnat* in heart could cause lipotoxic cardiomyopath, it remained unknown whether cardiac *Nmnat* gain of function could improve heart function and increase the heart’s resistance to lipotoxic cardiomyopath. So, to further confirm that cardiac *Nmnat* and cardiac Nmnat/NAD^+^/SIR2 pathways were key regulators of heart defects, cardiac *Nmnat* overexpression was built by UAS/hand-Gal4 in flies, and then these flies were fed a HFD.

The results showed that cardiac relative *Nmnat* expression in Nmnat-OE group flies was higher than Nmnat-C group flies (LSD test, P < 0.01), which suggested that the cardiac *Nmnat* overexpression was successfully constructed by UAS/hand-Gla4 in flies(**Fig.6-a**). Besides, for cardiac Nmnat/NAD^+^/SIR2 pathways, the results showed that the cardiac NAD^+^ level, *SIR2* expression, FOXO expression, SOD activity level, and *PGC-1α* expression in Nmnat-OE group flies were lower than Nmnat-C group flies (LSD test, P < 0.05 or P < 0.01) (**Fig.6-b, c, d, e, and g**), but the MDA level in Nmnat-OE group flies was higher than Nmnat-C group flies (LSD test, P < 0.05) (**Fig.6-f**). These results suggested that overexpression of cardiac *Nmnat* increased the activity of cardiac Nmnat/NAD^+^/SIR2 pathways and reduced the risk of oxidative damage to myocardial cells. Moreover, the cardiac TAG level in Nmnat-OE group flies was lower than Nmnat-C group flies (LSD test, P < 0.01), and the cardiac *bmm* expression in Nmnat-OE group flies was higher than Nmnat-C group flies (**Fig.6-h and i**). These results suggested that cardiac *Nmnat* overexpression reduced heart lipid accumulation. Finally, the heart rate, and fibrillation in Nmnat-OE group flies were lower than Nmnat-C group flies (LSD test, P < 0.05), and the fractional shortening, diastolic diameter, and systolic diameter in Nmnat-OE group flies were higher than Nmnat-C group flies (LSD test, P < 0.05 or P < 0.01) (**Fig.6-j, k, l, m, n, and o**). These results indicated that the cardiac *Nmnat* overexpression enhanced the heart contractility and reduced cardiac arrhythmia. Therefore, we claimed that cardiac *Nmnat* overexpression could improve heart function by increasing cardiac Nmnat/NAD^+^/SIR2 pathways activity.

Although previous results showed that cardiac *Nmnat* overexpression increased cardiac Nmnat/NAD^+^/ SIR2 pathways activity, decreased heart fat accumulation, strengthened heart contractility, and reduced fibrillation, it remained unclear whether this improvement of heart function induced by cardiac *Nmnat* overexpression could resist HFD-induced lipotoxic cardiomyopath. To further explore whether cardiac *Nmnat* overexpression could resist cardiac malfunction induced by a HFD, cardiac *Nmnat* overexpression flies were fed a HFD. The results displayed that there was no significant difference in cardiac NAD^+^ level, *SIR2* expression, FOXO expression, SOD activity level, *PGC-1α* expression, TAG level, *bmm* expression, heart rate, fractional shortening, diastolic diameter, systolic diameter, and fibrillation between Nmnat-OE flies Nmnat-OE+HFD flies (LSD test, P > 0.05) (**Fig.6-b, c, d, e, f, g, h, i, j, k, and o**), but the relative *Nmnat* expression in Nmnat-OE+HFD group flies were lower than Nmnat-OE group flies (LSD test, P < 0.05) (**Fig.6-a**). Besides, the results also showed that the cardiac relative *Nmnat* expression, the cardiac NAD^+^ level, *SIR2* expression, FOXO expression, SOD activity level, and *PGC-1α* expression in Nmnat-OE+HFD group flies were higher than Nmnat-C group flies (LSD test, P < 0.05 or P < 0.01) (**Fig.6-a, b, c, d, e, and g**), but the MDA level in Nmnat-OE+HFD group flies was lower than Nmnat-C group flies (LSD test, P < 0.05) (**Fig.5-f**). The cardiac TAG level in Nmnat-OE+HFD group flies was lower than Nmnat-C group flies (LSD test, P < 0.01), and the cardiac *bmm* expression in Nmnat-OE+HFD group flies was higher than Nmnat-C group flies (**Fig.6-h and i**). The heart rate and fibrillation in Nmnat-HFD group flies were lower than Nmnat-C group flies (LSD test, P < 0.05), and the fractional shortening, diastolic diameter, and systolic diameter in Nmnat-HFD+OE group flies were higher than Nmnat-C group flies (LSD test, P < 0.05 or P < 0.01) (**Fig.6-j, k, l, m, n, and o**). These results indicated that cardiac *Nmnat* overexpression could resist lipotoxic cardiomyopath induced by a HFD by activating cardiac Nmnat/NAD^+^/SIR2 pathways.

### The impact of cardiac *Nmnat* gene on the climbing ability and longevity

It has been reported that the *Nmnat* overexpression of whole body can improve health-span by enhancing stress resistance and locomotor activity in aging *Drosophila* ^24^. Besides, our previous study has shown that a HFD causes a decrease in lifespan and mobility, but endurance exercise could protect the lifespan and mobility from a HFD ^4^. However, it remains unclear that the impact of cardiac *Nmnat* expression on the climbing ability and longevity of flies, the impact of endurance exercise on the climbing ability and longevity of cardiac *Nmnat* knock-down flies, and the impact of a HFD on the climbing ability and longevity of cardiac *Nmnat* expression flies.

The results displayed that either cardiac *Nmnat* overexpression or *Nmnat* knock-down could not significantly affect the climbing index in young or adult flies (LSD test, P > 0.05) (**Fig.7-a and b**). However, the results showed that the relative climbing index in 7-week and *Nmnat*-KD group flies was lower than 7-week and *Nmnat-C* group flies (LSD test, P < 0.05), and the relative climbing index in 5- or 7-week and *Nmnat*-KD+E group flies was higher than 5- or 7-week and *Nmnat*-KD group flies and *Nmnat-C* group flies (LSD test, P < 0.05 or P < 0.01) (**Fig.7-c**). Besides, the results showed that the relative climbing index in 5-week and *Nmnat-OE* group flies was higher than 5-week and *Nmnat-C* group flies (LSD test, P < 0.05), and the relative climbing index in 5- or 7-week and *Nmnat*-OE+HFD group flies was lower than 5- or 7-week and *Nmnat*-OE group flies and *Nmnat-C* group flies (LSD test, P < 0.01) (**Fig.7-d**). For the lifespan, the results displayed that the lifespan of cardiac *Nmnat*-KD group flies was shorter than cardiac *Nmnat-C* group flies (a log-rank test, P < 0.05), and the lifespan of cardiac *Nmnat*-KD+E group flies was longer than cardiac *Nmnat*-KD group flies and cardiac *Nmnat-C* group flies (a log-rank test, P < 0.05 or P < 0.01) (**Fig.7-e**). Moreover, the results displayed that the lifespan of cardiac *Nmnat*-OE group flies was longer than cardiac *Nmnat-C* group flies (a log-rank test, P < 0.01), and the lifespan of cardiac *Nmnat*-OE+HFD group flies was shorter than cardiac *Nmnat*-OE group flies and cardiac *Nmnat-C* group flies (a log-rank test, P < 0.01) (**Fig.7-f**). These results suggested that cardiac *Nmnat* knock-down reduced the climbing ability of older flies and lifespan, but endurance exercise could completely reverse the adverse effect of climbing ability and lifespan induced by cardiac *Nmnat* knock-down. Cardiac *Nmnat* overexpression increased the climbing ability of older flies and lifespan, but it could not resist the adverse the effect of climbing ability and lifespan induced by a HFD.

**Fig.7.**
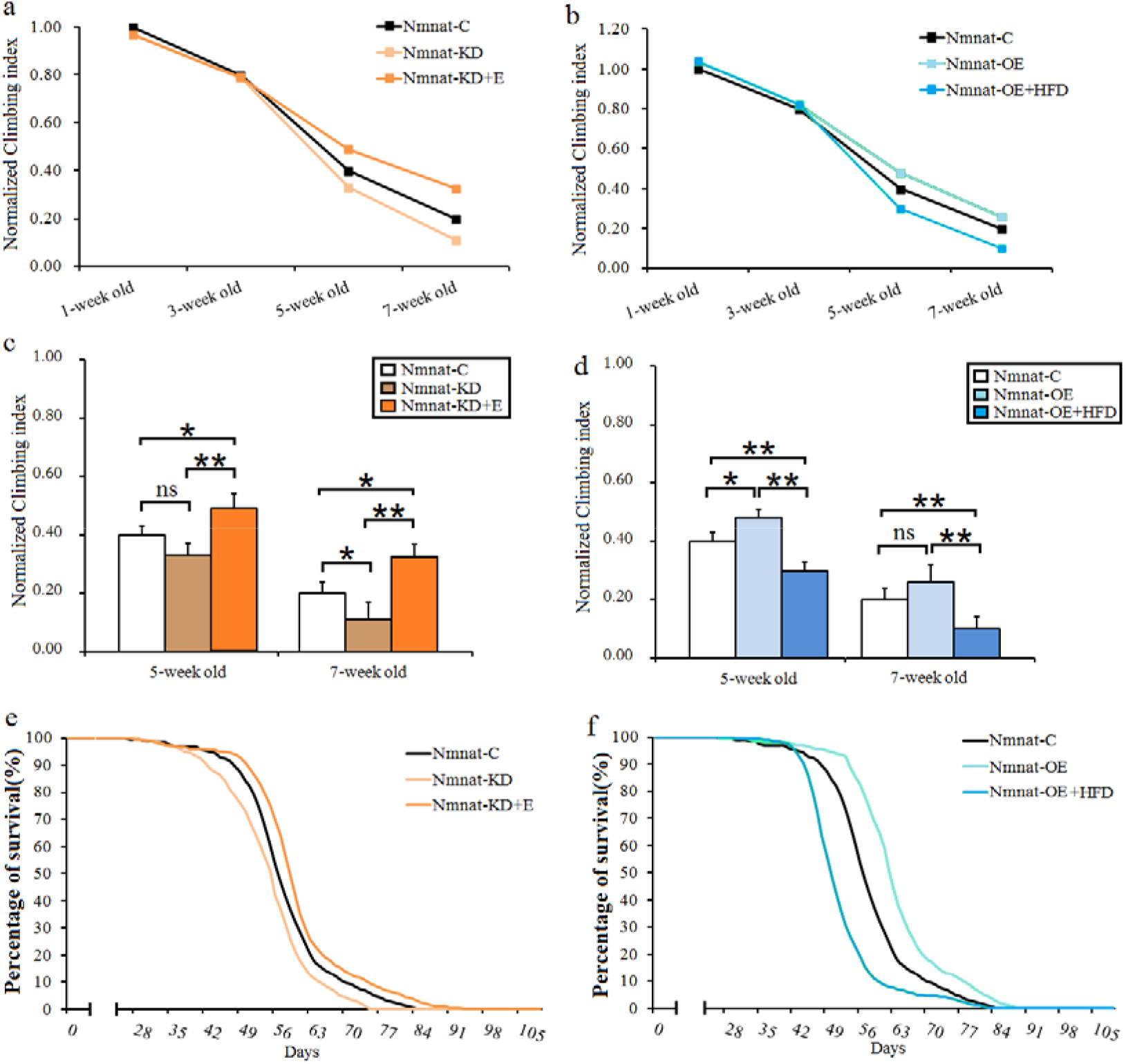
The effect of cardiac *Nmnat,* exercise, and HFD on mobility and lifespan. **(a)** Relative climbing index changes with ages **in** cardiac *Nmnat* knock-down flies. **(b)** Relative climbing index changes with ages in cardiac *Nmnat* overexpression flies. **(c)** Relative climbing index in cardiac *Nmnat* knock-down flies. **(d)** Relative climbing index in cardiac *Nmnat* overexpression flies. **(e)** The cur**ves** of the percentage of survival in cardiac *Nmnat* knock-down flies. **(f)** The curves of the percentage of survival in cardiac *Nmnat* overexpression flies. The sample size of climbing index was 100-120 flies for each group, and data are represented as means ± SEM. *P <0.05; **P <0.01.The sample size of lifespan was 200-220 flies for each group, and using a non-parametric followed by a log-rank test for analyze lifespan.

## Discussion

In both mammals and fruit flies, a HFD causes obesity and cardiac defects. For example, A HFD causes cardiac lipid accumulation, cardiac contractility debility, fractional shortening reduction, conduction blocks, and severe structural pathologies ^72^ī Besides, a HFD can induce expression change of some key genes, such as it can decrease heart *bmm, FOXO,* and *PGC-1α* expression and increase heart *TOR* and *FAS* expression^5,8^. In this study, the results again confirmed previous researches. In addition, we also found that a HFD reduced cardiac Nmnat/NAD^+^/SIR2 pathways activity, such as it reduced cardiac *Nmnat* expression, NAD^+^ level, *SIR2* expression, *FOXO* expression, and *PGC-1α* expression. Since over-expression of Sir2 deacetylase or increasing NAD^+^ levels activates transcriptional activity of PGC-1α and FOXO^26,27^, the cardiac Nmnat/NAD^+^/SIR2/PGC-1α and Nmnat/NAD^+^/SIR2/FOXO activity were suppressed by a HFD. The FOXO is an upstream regulator of SOD, and SOD is an oxygen free radical scavenging enzyme. So, a HFD may increase oxidative damage to cardiac cells by decreasing Nmnat/NAD^+^/SIR2/FOXO/SOD activity and increasing MDA level. Moreover, deacetylation of PGC-1α has been shown by several studies to be dependent on Sirt1 and NAD^+^ activity, which increases PGC-1α’s transcriptional activity^28,29^. PGC-1α is a key transcriptional regulator of mitochondrial biogenesis and function, and decreasing PGC-1α expression causes mitochondrial loss and defective mitochondrial function^30^. So, a HFD decreased cardiac *PGC-1α* expression by decreasing Nmnat/NAD^+^/SIR2 activity, which led to a decreased mitochondrial content in myocardial cells. Accumulating study indicates that HFD-induced obesity decreases the levels of NAD^+^ by several ways. For example, HFD-induced an increased oxidative stress reduces NAD^+^ level via PARP-1 activation mediated cell death ^31–35^ So, a HFD caused a severe lipid accumulation in heart. At the same time, a HFD reduce cardiac Nmnat/NAD^+^/SIR2 pathways by reducing NAD^+^ level, which caused more severe oxidative stress and mitochondrial dysfunction. Changes in these molecular pathways by a HFD led to decreased contractility in cardiac myocytes.

Although our previous results indicated that exercise resistance to HFD-induced lipotoxic cardiomyopath is associated with activation of cardiac Nmnat/NAD^+^/SIR2 pathways, it remained unclear whether the cardiac cardiac Nmnat/NAD^+^/SIR2 pathways could regulate the formation of HFD-induced heart defects. To confirm this, the caridac *Nmnat* gene loss-of-function and gain-of-function were built by UAS/hand-Gal4 system. In the heart of fly, accumulating evidence has confirmed that some key genes loss of function in heart causes cardiac dysfunction and lipid accumulation, such as cardiac *bmm, PGC-1α,* and *FOXO,* which was similar to lipotoxic cardiomyopath. On the contrary, these genes gain-of-function protects heart from HFD HFD-induced lipotoxic cardiomyopath ^5,8^. So, cardiac *PGC-1α, FOXO,* and *bmm* has been identified are key antagonists of HFD-induced heart defects in flies. These studies also suggested that making the gene gain- or loss-of-function has become a classic method to confirm gene function. It has been reported that the *Nmnat* gene loss-of-function in brain causes axons degeneration, Wallerian degeneration, and neurodegeneration, and these diseases can be prevented by the *Nmnat* gene gain-of-function ^36–38^. In this study, we found that knock-down of heart *Nmnat* decreased cardiac Nmnat/NAD^+^/SIR2 pathways activity and SOD activity, increased cardiac fat accumulation and MDA level, reduced cardiac contractility, and elevated cardiac fibrillation. These changes of heart induced by cardiac *Nmnat* knock-down were similar to HFD-induced lipotoxic cardiomyopath. Since the Nmnat reversibly catalyzes the important step in the biosynthesis of NAD from ATP and NMN ^19^, loss of cardiac Nmnat decreased heart NAD^+^ level, and eventually caused lipotoxic cardiomyopath via inhibiting cardiac Nmnat/NAD^+^/SIR2/PGC-1α and Nmnat/NAD^+^/SIR2/FOXO activity ^39–41^. Inversely, overexpression of heart *Nmnat* up regulated cardiac Nmnat/NAD^+^/SIR2 pathways activity and SOD activity, decreased cardiac fat accumulation and MDA level, enhanced cardiac contractility, and reduced cardiac fibrillation. These results suggested that activation of cardiac Nmnat/NAD^+^/SIR2 pathways reduced the incidence rate of lipotoxic cardiomyopath. However, we did not know yet whether overexpression of heart *Nmnat* could effectively resist HFD-induced lipotoxic cardiomyopath. To further identify this, the heart *Nmnat* overexpression flies were fed a HFD. The results showed that a HFD could not reduce cardiac Nmnat/NAD^+^/SIR2 pathways activity and SOD activity, and it couldn’t increase cardiac fat accumulation and MDA level, decrease cardiac contractility, and elevated cardiac fibrillation in cardiac *Nmnat* overexpression flies. Therefore, we claimed that the cardiac Nmnat/NAD^+^/SIR2 pathways were important antagonists of HFD-induced heart defects in flies.

Many studies have confirmed that appropriate endurance exercise is a healthy and economical way to prevent and cure obesity, and it is also considered a good way to improve heart functional in obesity or old individuals^42^. For instance, exercise training strengthens the heart’s ability to use fatty acids to provide energy by increasing the activity of related enzymes, which prevents lipid excessive accumulation in heart^43^. Besides, exercise training improves heart function such as cardiac contractibility and reduces heart failure in obesity individuals ^44–47^. Moreover, it has been reported that exercise increases cardiac SOD and FOXO activity and enhances the ability of myocardial cells to resist oxidative stress ^12,48^. Finally, although exercise increases muscle NAD^+^ levels and neurons NAD^+^ levels, which activates transcriptional activity of *PGC-1α* and increases mitochondrial density in muscles and neurons ^26,27^, it remains unclear whether exercise can prevent HFD-induced lipotoxic cardiomyopath via up regulating cardiac Nmnat/NAD^+^/SIR2 pathways activity. In this study, we found endurance exercise prevented HFD-induced cardiac fat accumulation, heart contractility reduction, and fibrillation increase, which are consistent with previous studies ‘Λ In addition, we found that endurance exercise resisted HFD-induced cardiac lipid accumulation by increasing *bmm* expression, which promoted the catabolism of fatty acids. Moreover, we found endurance exercise could prevent cardiac Nmnat/NAD^+^/SIR2 pathways inhibition induced by a HFD, which may be the molecular regulation mechansim of exercise resistance to HFD-induced lipotoxic cardiomyopath. To further confirm this hypothesis, the cardiac *Nmnat* knock-down flies were taken endurance exercise. We found endurance exercise improved lipotoxic cardiomyopath induced by cardiac *Nmnat* knock-down by up-regulating cardiac Nmnat/NAD^+^/SIR2 pathways activity. So, we claimed that activation of cardiac Nmnat/NAD^+^/SIR2 pathways were the key molecular mechanism of endurance exercise resistance against heart defects.

Besides, we also explore the effects of cardiac *Nmnat* gene, endurance exercise, and a HFD on the mobility and lifespan of fruit fly. In humans, Heart disease is a major cause of death in elderly individuals^49^. As we knew, heart function is an important factor in determining exercise ability since it is closely related to the rate of oxygen transport in the cells and tissues ^50,51^. It has been reported that the *Nmnat* overexpression of whole body can improve health-span by enhancing stress resistance and locomotor activity in aging *Drosophila* ^24^. In this study, we found knock-down of cardiac Nmnat induced locomotor impairment of older flies, and it notably reduced lifespan. These results indicated that knockdown of cardiac Nmnat reduced health-span of flies. In youth and adulthood, the heart shows a strong ability to contract ^52,53^. So, the cardiac Nmnat knock-down may not significantly reduce mobility by decreasing oxygen transport. However, cardiac *Nmnat* knock-down accelerated heart dysfunction with aging since NAD^+^ was an important factor to regulate cellular senescence^54,55^, and this eventually reduced climbing ability of flies. Our results also showed that endurance exercise improved climbing ability and extent their life-span in cardiac *Nmnat* knock-down flies, and the climbing ability and lifespan in cardiac *Nmnat* knock-down and trained flies were better than cardiac *Nmnat* normal flies. The reason may be that exercise improved not only heart function, but also skeletal muscle, brain, and other vital organs. In addition, we found cardiac *Nmnat* overexpression increased the climbing ability of older flies and lifespan, but it could not resist the adverse the effect of climbing ability and lifespan induced by a HFD. It has been reported that a HFD causes obesity in both animals and flies, which increases the incidence of diabetes, fatty liver, stroke, and hypertension ^56–58^. This may be the reason that cardiac *Nmnat* overexpression could not prevent HFD-induced locomotor impairment and lifespan reduction.

In summary, current results confirmed that cardiac *Nmnat* gene plays an important role in regulating the formation of heart defects. Overexpression of cardiac *Nmnat* activated cardiac Nmnat/NAD^+^/SIR2 pathways, which resisted HFD-induced lipid accumulation, cardiac dysfunction, and oxidative stress, but it could not reverse HFD-induced lifespan shortening and locomotor impairment of older flies. On the contrary, a HFD or cardiac *Nmnat* knock-down had similar effects on the heart, inducing lipid accumulation, cardiac dysfunction, and oxidative stress. However, endurance exercise resisted HFD- or *Nmnat* knock-down-induced heart defects via activating cardiac *Nmnat*/NAD^+^/*SIR2* pathways **(Fig.8)**.

**Fig.8.**
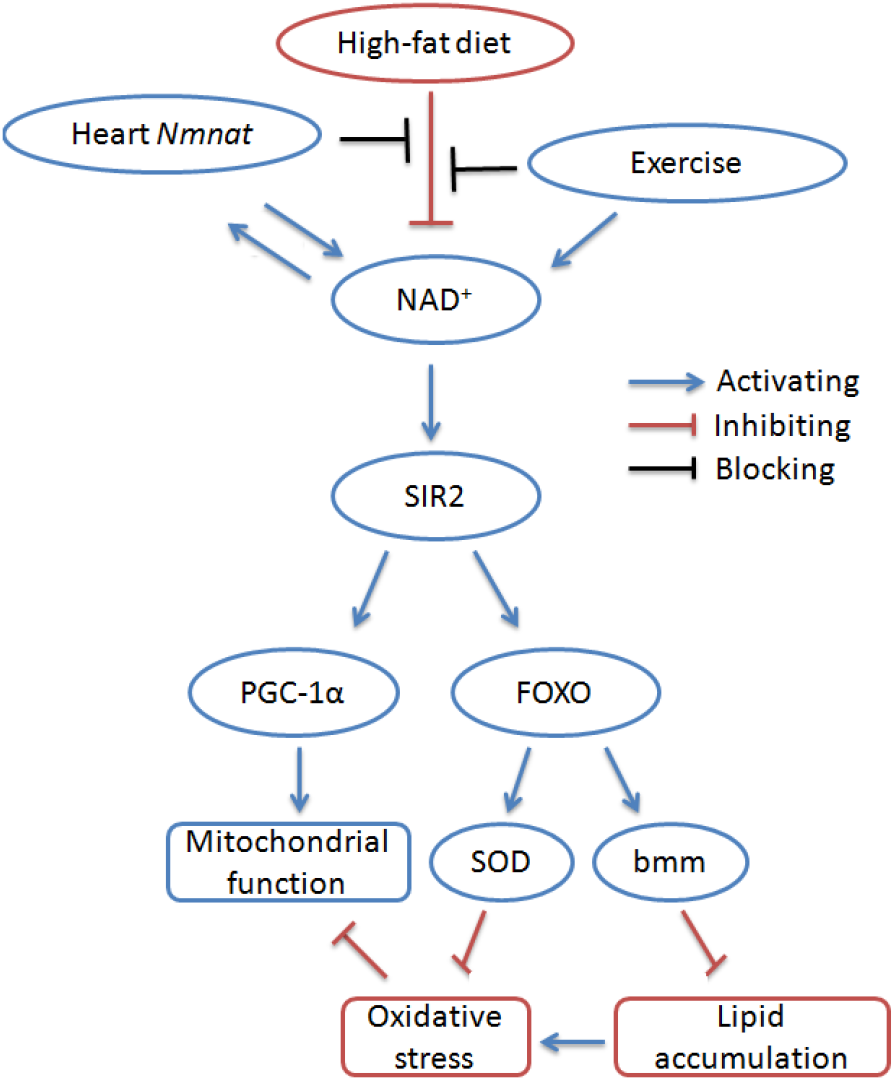
Current results confirmed that cardiac *Nmnat* gene plays an important role in regulating the formation of heart defects. Overexpression of cardiac *Nmnat* activated cardiac Nmnat/NAD^+^/SIR2 pathways, which resisted HFD-induced lipid accumulation, cardiac dysfunction, and oxidative stress. On the contrary, a HFD or cardiac *Nmnat* knock-down had similar effects on the heart, inducing lipid accumulation, cardiac dysfunction, and oxidative stress. However, endurance exercise resisted HFD- or *Nmnat* knock-down-induced heart defects via activating cardiac *Nmnat*/NAD^+^/*SIR2* pathways.

## Materials and methods

### Fly stocks, diet and husbandry

The *w^1118^* and *hand-Gal4* flies were a gift from Xiu-shan Wu (Heart Development Center of Hunan Normal University). UAS-*Nmnat*-overexpression *(y^1^ w*; P{UAS-Nmnat.Z}2/CyO)* flies were obtained from the Bloomington Stock Center. UAS-*Nmnat*-RNAi *(P{KK101988}VIE-260B)* flies were obtained from the Vienna Drosophila RNAi Center. To build different expression of the *Nmnat* gene in fly heart, male *hand-Gal4* flies were crossed to female *w^1118^* flies, UAS-*Nmnat*-overexpression flies, and UAS-*Nmnat*-RNAi flies. The Gal4/upstream activating sequence (UAS) system was one of the most powerful tools for targeted gene expression. It was based on the properties of the yeast GAL4 transcription factor which activates transcription of its target genes by binding to UAS *cis*-regulatory sites. In *Drosophila,* the two components are carried in separate lines allowing for numerous combinatorial possibilities^59^. The bipartite system is commonly used in gain-of-function analysis, and by combining with the RNA interference technology, it can also be applied in loss-of-function analysis^60^. The *“hand-Gal4>w^1118^”, “hand-Gal4>UAS-Nmnat-overexpressiotì\* and “*hand-Gal4>*UAS-*Nmnat*-RNAi” were represented as “*Nmna-*Control (*Nmnat*-C)”, “*Nmnat*-overexpression(*Nmnat*-OE)”, and “*Nmnat*-knock-down(*Nmnat*-KD)”. All UAS and GAL4 insertions were backcrossed into the *w^1118^* line at least 10 times to avoid excess phenotypes affecting the experimental results. All group flies were raised to the fourth weekend, and then flies were trained in their five weeks old since we found flies were more sensitive to exercise in this time.

Normal food contained 10% yeast, 10% sucrose and 2% agar. The high fat diet was made by mixing 30% coconut oil with the food in a weight to volume ratio with the normal food ^5^. All HFD group flies were fed a HFD from 21-day old and were exposed to the HFD for 2 weeks. During the experimental time course, flies were housed in a 22±1°C incubator with 50% humidity and a 12-h light/dark cycle. This environment could keep the coconut oil food in solid state since the melting point of coconut oil is about 24°C, thus ensuring that flies would not get stuck in the oily food. Fresh food was provided every other day for the duration of the experiment. All group flies were raised to the fourth weekend. Flies were trained or fed a HFD at their five weeks old since we found flies were very sensitive to exercise or high-fat diet at this time.

### Exercise training device and protocols

When constructing the exercise device, the advantage of the flies’ natural negative geotaxis behavior was taken to induce upward walking. All exercise groups flies started exercise from when they were 21-day old, and underwent a 2-week-long exercise program. Vials, with the diet housing 25 flies each, were loaded horizontally into a steel tube that was rotated about its horizontal axis by an electric motor, with a gear regulating its shaft speed. Thus, with the accompanying rotating steel tube, each vial was rotated along its long axis, which made the flies climb (The Tread Wheel)^61^. Most flies continued to respond by climbing throughout the exercise period. The few that failed to climb were actively walking at the inner wall of the vial ^15,61,62^. Flies were exercised in vials with a 2.8-cm inner diameter, rotated at 0.16 rev/s. Flies were exercised for 1.5 hours.

### Semi-intact Drosophila preparation and image analysis

Flies were anesthetized with FlyNap for 2-3 min. And the head, ventral thorax, and ventral abdominal cuticle were removed by special glass needles to expose the heart and abdomen. Dissections were done under oxygenated artificial hemolymph. These semi-intact preparations were allowed to equilibrate with oxygenation for 15-20 min before filming. Image analysis of heart contractions was performed using high-speed videos of the preparations. Videos were taken 120-130 frames per second using a Hamamatsu (McBain Instruments, Chats worth, CA) EM-CCD digital camera on a Leica (McBain Instruments, Chatsworth, CA) DM LFSA microscope with a 10 immersionlens. To get a random sampling of heart function, a single 30-s recording was made for each fly. All images were acquired and contrast enhanced by using Simple PCI imaging software (Compix, Sewickley, PA). The heart physiology of the flies was assessed using a semiautomated optical heartbeat analysis program that quantifies heart rate, fractional shortening, diastolic diameter, systolic diameter, and fibrillation^63^. The sample size was 30 flies for each group.

### ELISA assay

The cardiac TAG, NAD^+^, MDA, and SOD levels were measured by ELISA assay (The insect TAG, NAD, MDA, and SOD ELISA Kits were provided by mlbio of Shanghai). Fly hearts were homogenized in PBS (PH7.2-7.4). Rapidly frozen with liquid nitrogen, maintain samples at 2-8°C after melting, Homogenized by Grinders, centrifugation 20-min at the speed of 2000-3000 r.p.m. remove supernatant. The steps were as the manufacturer’s instructions. Every factor measurement needed repeated 3 times, and each group required total 240 hearts in ELISA assay. The normal lines of the TAG, NAD^+^, MDA, and SOD were worked out to calculate heart TAG levels, NAD^+^ levels, MDA, and SOD levels.

### qRT-PCR

About 80 hearts of each group were homogenized in Trizol. Firstly, 10 μg of the total RNA was purified by organic solvent extraction from the Trizol (TRIzol, Invitrogen). The purified RNA was treated with DNase I (RNase-free, Roche) and used to produce oligo dT-primed cDNAs (SuperScript II RT, Invitrogen), which were then used as templates for quantitative real-time PCR. The rp49 gene was used as an internal reference for normalizing the quantity of total RNAs. Real-time PCR was performed with SYBR green using an ABI7300 Real time PCR Instrument (Applied Biosystems). Expression of the various genes was determined by the comparative CT method (ABI Prism 7700 Sequence Detection System User Bulletin #2, Applied Biosystems). Primer sequences of *PGC-p/_* were as follows: F: 5′-TGTTGCTGCT ACTGCTGCTT-3′ R:5′-GCCTCTG CATCACCTACACA-3′. Primer sequences of *bmm* were as follows: F: F: 5′-ACTGCACATTTCGCTTACCC-3′ R: 5′-GAGAATCCGGGTATGAAGCA-3′. Primer sequences of *SIRT1* were as follows: F: 5′-GCCCAAGAACAACATAACAAGC-3′ R: 5′-CGAGATGATGCCACCTACCAC-3′. Primer sequences of *FOXO* were as follows: F: 5′-AACAACAGCAGCATCAGCAG-3′ R: 5′-CTGAACCCGAGCATT CAGAT-3′. Primer sequences of *Rp49* were as follows: F: 5′-CTAAGCTGTCGCACAAATGG-3′ R:5′-AACTTCTT GAATCCGG TGGG-3′.

### Negative geotaxis assay

The climbing apparatus consisted of an 18-cm-long vial with an inner diameter of 2.8 cm, and flies were allowed to adapt to the vial for ten minutes before assessing negative geotaxis. Sponges were placed in the ends of the tube to prevent escape while allowing air exchange. With a light box behind the vials, the rack was tapped down five times and on the fifth, a timed digital camera snapped a picture after 8 seconds. The extent of climbing could be analyzed visually or by imaging software. Five pictures of each group were taken and averaged to arrive at a fixed score for each vial. The total score for all the flies in a vial was tallied, and then divided by the number of flies in the vial to generate the “Climbing Index” for that trial. Each vial was subjected to 5 trials, and then the indexes from the five trials were averaged.

### Lifespan assays

Dead flies were recorded daily. Lifespan was estimated for each fly as number of days alive from day of eclosion to day of death. Mean and median lifespan and survival curves were primarily used to characterize lifespan. Sample sizes were 200-220 flies per group.

### Statistical analyses

A 2-way ANOVA was used to analyze the effects of HFD and exercise on the heart in Nmnat-C flies. The 1-way analysis of variance (ANOVA) with least significant difference (LSD) tests was used to identify differences among the “Nmnat-C group”, “Nmnat-KD group”, and “Nmnat-KD+E group”. The 1-way analysis of variance (ANOVA) with least significant difference (LSD) tests was used to identify differences among the “Nmnat-C group”, “Nmnat-OE group”, and “Nmnat-OE+HFD group”. Analyses were performed using the Statistical Package for the Social Sciences (SPSS) version 16.0 for Windows (SPSS Inc., Chicago, USA), with statistical significance set at *P<0.05.* Data are represented as means ± SEM.

## Acknowledgements

The authors thank Xiu-shan Wu (The Center for Heart Development, College of Life Science, Hunan normal University) for supporting *Drosophila* of *w^1118^, hand-Gal4*, and heart Shoot software technology. We also thank Karen Ocorr and Rolf Bodmer (American burnham medical institute of neurology and aging center) for supporting semi-automatic optical echocardiography analysis software.

## Author Contributions

Research idea and study design: D.t.W., L.Z.; data acquisition: D.t.W.; data analysis/interpretation: Y.F., D.t.W., K.L.; statistical analysis: D.t.W.,; supervision: L.Z., H.w.Q. Each author contributed during manuscript drafting or revision and approved the final version of the manuscript.

## Associated Data

### Data Availability Statement

All the generated data and the analysis developed in this study are included in this article.

